# 2’-Fucosyllactose helps butyrate producers outgrow competitors in infant gut microbiota simulations

**DOI:** 10.1101/2023.03.10.532059

**Authors:** David M. Versluis, Ruud Schoemaker, Ellen Looijesteijn, Jan M. W. Geurts, Roeland M. H. Merks

## Abstract

A reduced capacity for butyrate production by the early infant gut microbiota is associated with negative health effects, such as inflammation and the development of allergies. Here we develop new hypotheses on the effect of the prebiotic galacto-oligosaccharides (GOS) or 2’-fucosyllactose (2’-FL) on butyrate production by the infant gut microbiota using a multiscale, spatiotemporal mathematical model of the infant gut. The model simulates a community of cross-feeding gut bacteria at metabolic detail. It represents the gut microbiome as a grid of bacterial populations that exchange intermediary metabolites, using 20 different subspecies-specific metabolic networks taken from the AGORA database. The simulations predict that both GOS and 2’-FL promote the growth of *Bifidobacterium*, whereas butyrate producing bacteria are only consistently abundant in the presence of propane-1,2-diol, a product of 2’-FL metabolism. The results suggest that in absence of prebiotics or in presence of only GOS, bacterial species, including *Cutibacterium acnes* and *Bacteroides vulgatus*, outcompete butyrate producers by feeding on intermediary metabolites. In presence of 2’-FL, however, production of propane-1,2-diol specifically supports butyrate producers.

## 1 Introduction

Infants develop a complex microbiota shortly after birth, which is important for healthy growth and development [70]. Here we focus on butyrate, a short-chain fatty acid (SCFA) that is produced in significant amounts by the gut bacteria [20] and is absorbed by the gut colonocytes. Production of butyrate by the microbiota has been suggested to improve the health of infants in a number of ways. Firstly, butyrate in the gut is a key energy source for the gut epithelium, making it important for maintaining the gut barrier function [58]. A breakdown of the gut barrier function due to a lack of butyrate is associated with diseases such as inflammatory bowel disease and rectal cancer [58, 82]. Butyrate production in young infants specifically is associated with a reduced risk of allergies and allergy-associated atopic eczema [11, 51, 81]. Infant butyrate producing bacteria provide protection against food allergies when transplanted into a mouse model [24], suggesting causality. Butyrate production is also associated with a reduced risk of colic in infants [17]. Butyrate also modulates the immune system throughout the body, inhibiting inflammation and carcinogenesis [33]. These data suggest it may be desirable to stimulate butyrate production in the infant gut. Using mechanistic computational modeling, here we investigate how stimulation of butyrate producing bacteria may be achieved in the early infant gut microbiota through supplementation with prebiotics.

Microbiota composition and metabolism are influenced by endogenous factors, e.g., gut maturity and inflammation, and exogenous factors, e.g., nutrition, probiotics, and antibiotics. Here we focus on nutrition, which is the primary exogenous factor. Human milk and many infant formulas contain prebiotics such as galacto-oligosaccharides (GOS) and 2’-fucosyllactose (2’-FL), which influence the composition of the gut microbiota and are associated with beneficial health effects for the infant, such as a decreased risk to require antibiotics [7] and reduced manifestation of allergies [49, 67, 28]. It has been hypothesized that some of the health effects associated with prebiotics may be linked to indirect stimulation of butyrate producing bacteria [73, 81]. Thus, both the capacity for butyrate production [11, 81], and prebiotics in nutrition by itself, particularly 2’-FL, have been linked to reduced manifestations of allergies [49, 67, 28]. Butyrate producing bacteria such as *Anaerobutyricum hallii* cannot directly consume GOS or 2’-FL, but they can consume metabolites of GOS or 2’-FL digestion [64]. The primary consumers of GOS and 2’-FL in the infant gut are *Bifidobacterium* spp.[8, 9]. Metabolites produced by *Bifidobacterium* spp., in turn, become important food sources for butyrate producing bacteria.

For example, *in vitro* it has been found that the butyrate producing bacterium *A. hallii* (formerly *Eubacterium hallii* [65]) can feed on lactate and propane-1,2-diol (1,2-PD), which are metabolites of *Bifidobacterium* spp. [64]. *A. hallii* can also coexist with *Bifidobacterium longum* ssp. *infantis in vitro* on a substrate of glucose or 2’-FL [64].

Despite these in vitro findings that demonstrate potential coexistence of *Bifidobacterium* spp. and butyrate producing bacteria, *in vivo*, i.e. in the infant gut microbiota, butyrate producing bacteria often only have a low abundance and butyrate is found in the feces of only 35% of infants [3]. It is unclear why butyrate producing bacteria and butyrate are not commonly abundant *in vivo*, given that *in vitro* cross-feeding on lactate occurs readily [64], and that lactate-producing *Bifidobacterium* species are abundant in the gut of most infants [4, 69]. Using computational modeling we explore the conditions that may stimulate butyrate producing bacteria *in vivo* in the infant gut. To this end we will compare simulations of simple microbial communities, such as those studied *in vitro*, with simulations of more complex communities that may more closely resemble the *in vivo* situation.

Briefly, the computational model suggests that in simple microbial communities, populations of butyrate producing bacteria can cross-feed on *Bifidobacterium* metabolites. However, in more complex communities the intermediary metabolites are consumed by competitors instead of butyrate producing bacteria. In the presence of 2’-FL, populations of butyrate producing bacteria are nevertheless supported. The mechanism suggested by our simulations is that *Bifidobacterium* produces 1,2-PD from 2’-FL, which specifically feeds butyrate producing species, allowing these to outgrow competing cross-feeders. We provide predictions for interactions in *in vivo* and *in vitro* systems and suggestions for *in vitro* verification of these predictions.

## 2 Results

### 2.1 Model outline

To develop new hypotheses on how oligosaccharides can stimulate the production of butyrate, we further develop a multiscale metabolic model (Fig. 1A & B) of the carbon metabolism of the infant gut microbiota [78]. The computational model is based upon our earlier models of the adult and infant microbiota [75, 78]. In comparison with these previous models, the present model simulates a larger number of small bacterial populations, using a larger, more diverse, and further curated set of metabolic models of gut bacteria from the AGORA database [44]. In particular, we have included the butyrate producers *A. hallii*, *Roseburia inulinivorans* and *Clostridium butyricum* and the digestion of the prebiotic oligosaccharides GOS and 2’-FL by *Bifidobacterium longum* ssp. *infantis*. The complete community model integrates these predictions of metabolism over space and time to create a multiscale model that covers the development and variation of the infant gut microbiota over the first three weeks of life. Other multiscale metabolic modelling techniques have been used previously to model the adult human microbiota in frameworks such as SteadyCom and Comets [12, 21]. The model presented here distinguishes itself from these frameworks by its focus on the infant gut microbiota, by including factors such as prebiotics and the initial presence of oxygen at birth.

**Figure 1:**
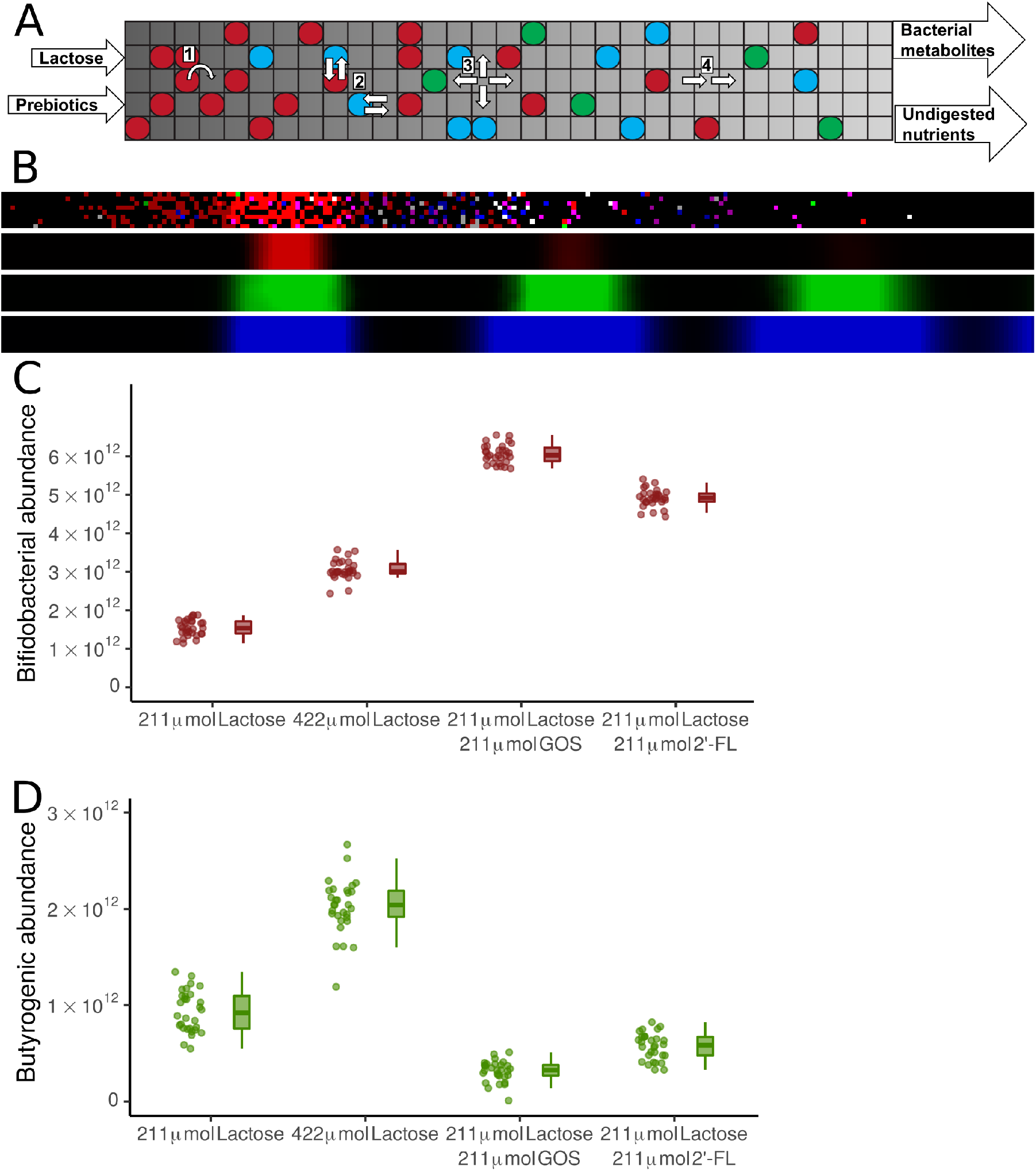
Model predicts coexistence of *Bifidobacterium* and butyrate producing bacteria in absence of competition (A) Schematic of the model. Circles represent bacterial populations, colour represents species. Flow through the tube is from left (proximal) to right (distal). Nutrients entered the system proximally. All metabolites leave the system distally. Lattice dimensions are schematic. (B) Screenshots of the model at a single time point, showing, from top to bottom, the bacterial layer, lactose, lactate, and acetate. Brightness indicates growth in the bacterial layer, and concentration in the metabolic layers. (C,D) Abundance of (C) *Bifidobacterium* spp., (D) butyrate producing bacteria, at the end of 21 days for 30 sets of simulations with no prebiotics, no prebiotics and additional lactose, with GOS, or with 2’-FL at the end of 21 days. n=30 for each condition, each simulation is represented by one dot.

Briefly, the spatial model simulates the ecology of an intestinal microbial ecosystem, and features genome-scale metabolic models (GEMs) of intestinal bacteria, spatial structuring, exchange of extracellular metabolites, and population dynamics. The system is simulated on a regular square lattice of 225 × 8 boxes of 2 × 2 mm, representing a typical infant colon of 45 × 1.6 cm. Each box contains a simulated metapopulation of one of a set of up to 20 of the most common bacterial species present in the infant gut [4] (Table 1), and concentrations of simulated nutrients and metabolites such as extracellular oligosaccharides and short-chain fatty acids. Based on the concentrations of metabolites, the systems predicts the growth rate for each metapopulation as well as the uptake and excretion rates of metabolites using a GEM taken from AGORA [43], a database of metabolic networks of intestinal bacteria. The system is initialised by distributing, on average, 540 populations over the system at random. Oxygen is introduced during initialisation, and water is always available.

**Table 1:** Species and subspecies included in the model. Colour indicates colour used in figures.

| Name | Phylum | Anaerobic status per [44] | Butyrate producing |
| --- | --- | --- | --- |
| *Bifidobacterium longum* ssp. *infantis* | Actinomycetota | Obligate anaerobe | no |
| *Bifidobacterium longum* ssp. *longum* | Actinomycetota | Obligate anaerobe | no |
| *Collinsella aerofaciens* | Actinomycetota | Obligate anaerobe | no |
| *Cutibacterium acnes* | Actinomycetota | Facultative anaerobe | no |
| *Rothia mucilaginosa* | Actinomycetota | Microaerophile | no |
| *Eggerthella* sp. *YY7918* | Actinomycetota | Nanaerobe | no |
| *Streptococcus oralis* | Bacillota | Facultative anaerobe | no |
| *Staphylococcus epidermidis* | Bacillota | Facultative anaerobe | no |
| *Gemella morbillorum* | Bacillota | Facultative anaerobe | no |
| *Enterococcus faecalis* | Bacillota | Facultative anaerobe | no |
| *Lactobacillus gasseri* | Bacillota | Facultative anaerobe | no |
| *Ruminococcus gnavus* | Bacillota | Obligate anaerobe | no |
| *Veillonella dispar* | Bacillota | Obligate anaerobe | no |
| *Anaerobutyricum hallii* | Bacillota | Obligate anaerobe | yes |
| *Roseburia inulinivorans* | Bacillota | Obligate anaerobe | yes |
| *Clostridium butyricum* | Bacillota | Obligate anaerobe | yes |
| *Parabacteroides distasonis* | Bacteroidota | Nanaerobe | no |
| *Bacteroides vulgatus* | Bacteroidota | Nanaerobe | no |
| *Haemophilus parainfluenzae* | Pseudomonadota | Aerobe | no |
| *Escherichia coli* SE11 | Pseudomonadota | Facultative anaerobe | no |

After initialisation, the model is simulated in timesteps representing three minutes of real time. Each timestep of the simulation proceeds as follows. Every 3 hours (i.e., 60 timesteps), a mixture of simulated lactose and/or oligosaccharides is added to the leftmost six columns of lattice sites. Then, each step, the model predicts the metabolism of each local population using flux balance analysis (FBA) based on the metabolites present in the local lattice site, the GEM of the species, and the enzymatic constraint. The enzymatic constraint limits the total amount of metabolism that can be performed by each local population per timestep by limiting the maximum summed flux for each FBA solution. The enzymatic constraint is determined by the local population size. This approach allows us to model metabolic switches and trade-offs

[45, 78]. The FBA solution includes a set of influx rates and efflux rates of metabolites that are used to update the environmental metabolite concentrations. The local populations are assumed to grow at a rate linearly proportional to the rate of ATP production[63], which is predicted by FBA by optimizing for ATP production rates. Populations may create a new population in a neighbouring lattice site if the local population is 200 times the initial size (Fig. 1A-1). Populations of more than 400 times the local size, which can only form when density if so high new populations cannot be created, stop metabolism to represent quiescence. Populations spread at random into adjacent lattice sites (Fig. 1A-2); metabolites diffuse and advect towards the back of the tube (Fig. 1A-3&4). To mimic excretion, metabolites and populations are deleted from the most distal column each timestep. To represent bacterial colonisation, new populations of randomly selected species are introduced into empty lattice sites at a small probability. All parameters are given in table 2. Details of the model are given in section Methods.

**Table 2:**
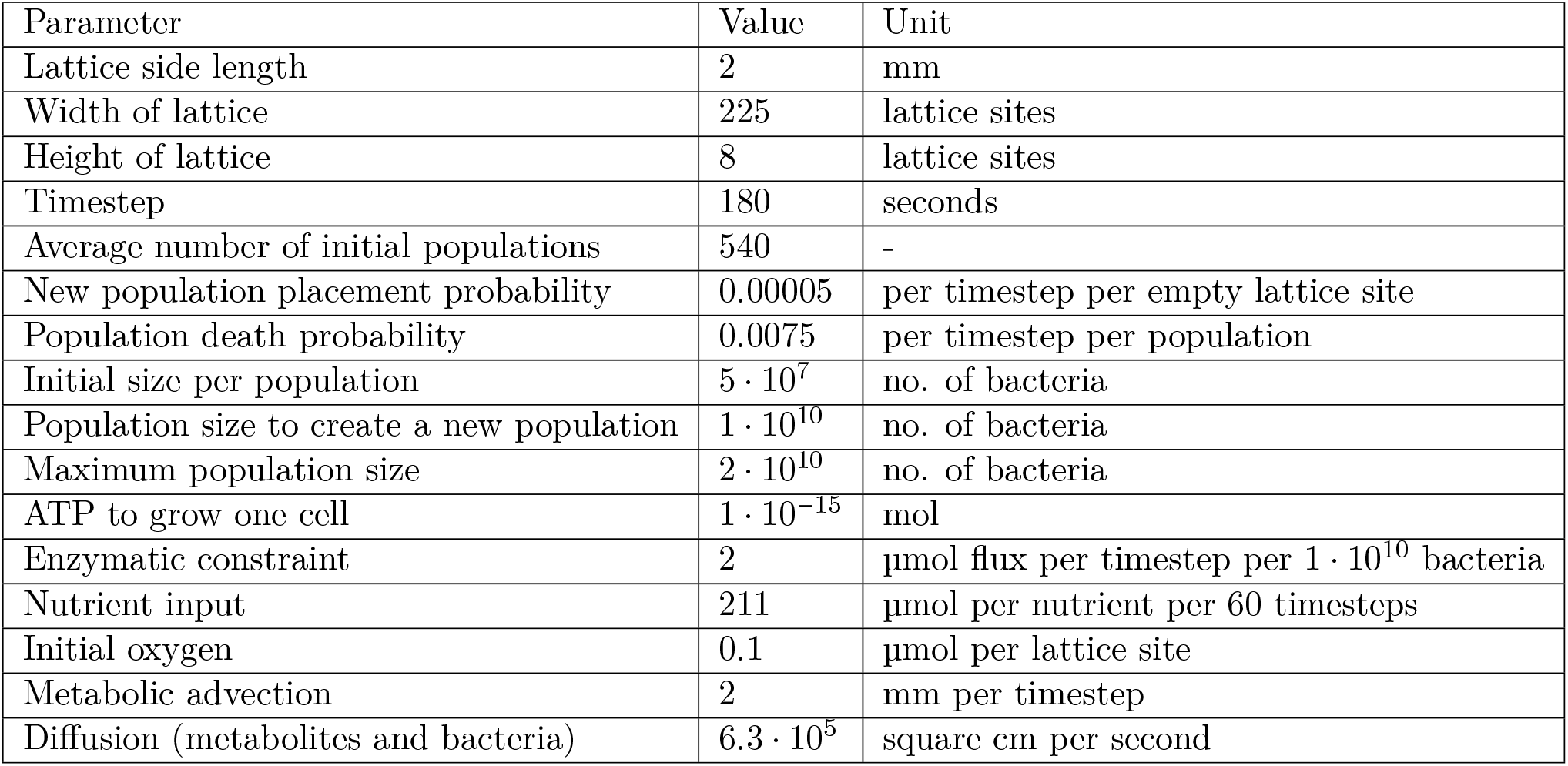
Parameters of the model

### 2.2 Model with simplified consortium of species predicts coexistence of butyrate producing bacteria and *Bifidobacterium*

We first simulated the model using a simplified consortium of species, the two *Bifidobacterium longum* subspecies (table 1) and three butyrate producing species: *Anaerobutyricum hallii*, *Clostridium butyricum*, and *Roseburia inulinivorans*. We performed 30 simulations for each of four conditions, in which the following sugars were introduced every three simulated hours: (1) 211 µmol lactose and no prebiotics, (2) 422 µmol lactose and no prebiotics, (3) 211 µmol lactose plus 211 µmol GOS, and (4) 211 µmol lactose plus 211 µmol 2’-FL. We estimated 211 µmol lactose to be a realistic amount of lactose to reach the infant colon, given infant intake and small intestinal uptake [5, 12]. As there is little absorption by the small intestine of prebiotics [27], the amount of prebiotics in the nutrition consumed by the infant would be much smaller than the amount of lactose. We also include the 422 µmol lactose condition to control for the possibility that effects in the conditions with prebiotics are due the larger amount of sugar present conditions, instead of their type. The condition with 422 µmol lactose does not correspond to an *in vivo* condition. We analyzed the abundance of each species at the end of 10080 timesteps, representing 21 simulated days. In each of the four conditions *Bifidobacterium* bacteria (Fig. 1C) and butyrate producing bacteria coexisted (Fig. 1D), and, paradoxically, butyrate producing bacteria were reduced in presence of prebiotics.

### 2.3 In presence of competitors, model predicts coexistence of butyrate producing bacteria and *Bifidobacterium* in the presence of 2’FL but not in presence of GOS

We next examined the behaviour of the system in the presence of a more complex consortium, consisting of all 20 species and subspecies listed in Table 1, simulating the same four conditions. In absence of prebiotics, regardless of the quantity of lactose, the model predicted that *Bifidobacterium*, *Bacteroides* and *Escherichia* became the most abundant genera after three weeks (Fig. 2A, S1 Video), consistent with *in vivo* observation [4, 69]. We also observed some abundance of Bacilli in accordance with *in vivo* observations [4, 19, 69]. The higher quantity of lactose resulted in a higher average abundance for all major groups. In absence of prebiotics, butyrate producing bacteria achieved a combined abundance over 1 · 10^10^ in only 4 of the 30 simulations with 211 µmol of lactose per 3 hours, and 6 of the 30 with 422 µmol of lactose (Fig. 2B). In the remaining simulations, the butyrate producing bacteria remained almost absent, staying below 1 · 10^10^ bacteria. In the simulations with GOS, *Bifidobacterium* was more abundant than in the condition without prebiotics (p<0.001,Fig. 2A) whereas the butyrate producing bacteria were not affected (p=0.18) (Fig. 2B). With GOS, butyrate producing bacteria also had a combined abundance of over 1 · 10^10^ bacteria at the end of 13 of the 30 simulations (Fig. 2B). Interestingly, in the condition with 2’-FL the abundance of butyrate producing bacteria was over 1 · 10^10^ bacteria at the end of 19 of 30 simulations (Fig. 2B), and the butyrate producing species were more abundant (Fig. 2A, S2 Video) than in the other conditions. Thus 2’-FL but not GOS stimulated butyrate producing bacteria in the complex community. To test for any concentration-dependence or cross-talk between 2’-FL and GOS we next performed sets of 30 simulations in presence of 211 µmol lactose and levels of 2’-FL and GOS varying between 21.1 µmol to 211 µmol per three hours and combinations thereof (Fig. S1). The amount of 2’-FL (p=0.017, Kruskal-Wallis rank sum test) but not that of GOS (p=0.658, Kruskal-Wallis rank sum test) affected the abundance of butyrate producing bacteria, further supporting the prediction that 2’-FL but not GOS stimulates butyrate producing bacteria in the complex community.

**Figure 2:**
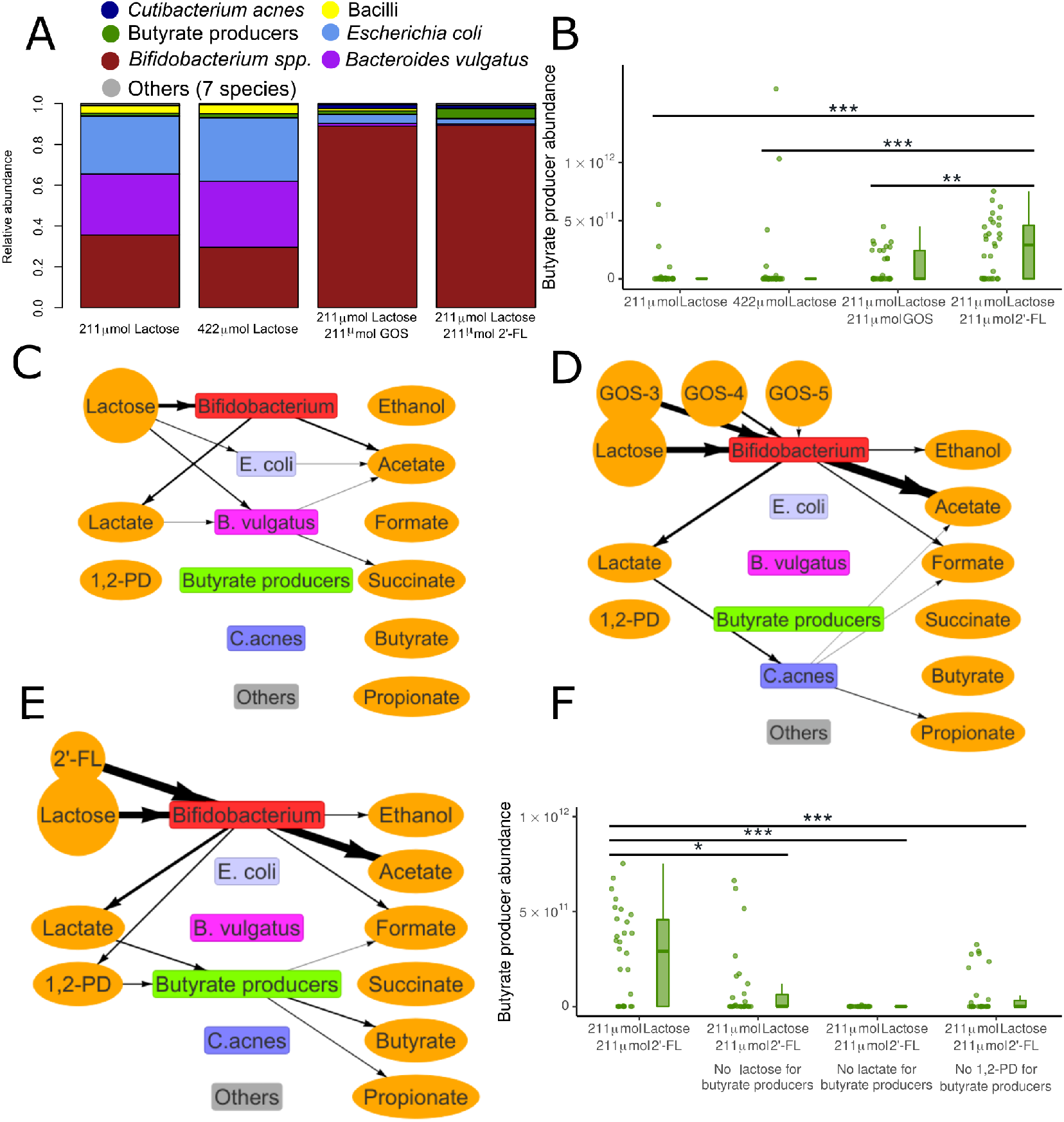
Unlike GOS, 2’-FL leads to stimulation of butyrate producing bacteria through 1,2-PD in the full simulated microbiota (A) Relative abundance of bacterial species in the condition with no prebiotics, no prebiotics and additional lactose, with GOS, or with 2’-FL at the end of 21 days. n=30 for each condition, each simulation is weighed equally. The key to the species in each group is in table 1. (B) Abundance of butyrate producing bacteria at the end of 21 days for the four conditions of A. n=30 for each condition. Each simulation is represented by one dot. p<0.001 for 2’-FL compared to no prebiotics and no prebiotics with additional lactose. p=0.004 for 2’-FL compared to GOS. (C,D,E) Visualisation of metabolic interactions in a sample simulation (C) without prebiotics (211 µmol lactose per three hours) (D) with GOS (DP3,DP4, and DP5 displayed separately) (E) with 2’-FL. Line width is scaled with the flux per metabolite over the last 60 timesteps, multiplied by the carbon content of the molecule, with a minimum threshold of 100 µmol atomic carbon. Data from last 3 hours, step 10020 to 10080. Circles indicate nutrients. (F) Abundance of butyrate producing bacteria with 2’FL at the end of 21 days. Uptake of lactose, lactate, or 1,2-PD by butyrate producing bacteria is disabled in the ‘no lactose’, ‘no lactate’, and ‘no 1,2-PD’ conditions respectively. p=0.010,p<0.001,p<0.001 for each disabled uptake compared to the baseline, respectively n=30 for each condition. Each simulation is represented by one dot. NS: Not significant, *: p<0.05, **:p<0.01, ***:p<0.001

In order to investigate why 2’-FL led to a more consistent abundance of butyrate producing bacteria we analysed the metabolic interactions between bacterial species. We visualised the network of metabolic fluxes between the bacteria using arrows between species and metabolite pools in Fig. 2C-E. The resulting diagrams show both primary consumption, i.e., uptake of nutrients such as lactose, GOS, and 2’-FL, and cross-feeding, i.e., uptake of metabolites produced by other species. Sample visualisations of the condition without prebiotics (211 µmol lactose) (Fig. 2C, S3 Video) and the condition with GOS (Fig. 2D) revealed co-occurrence of species and cross-feeding, but no butyrate production. In these simulations the cross-feeding metabolite lactate, which is a known substrate for butyrate producing bacteria [64], was consumed by *Bacteroides vulgatus* and *Cutibacterium acnes*, respectively. Butyrate formation only occurred in the sample simulation with 2’-FL (Fig. 2E). Only in presence of 2’-FL and not in the other conditions, was a flux of 1,2-PD directed towards the butyrate producing species (Fig. 2E and S4 Video). We therefore hypothesised that butyrate producing species may be more abundant in the model simulations with 2’-FL, because 2’-FL digestion by *Bifidobacterium* produces 1,2-PD as a cross-feeding substrate. 1,2-PD is a known *Bifidobacterium* metabolite from 2’-FL *in vitro* [64]. To test this hypothesis, we performed new sets of simulations with 2’-FL in which we blocked the uptake by butyrate producing bacteria of either lactose, lactate, or 1,2-PD, i.e, the uptake of metabolites most consumed by butyrate producing bacteria was disabled. Indeed, blocking the uptake of any of these metabolites led to a reduction of butyrate producing bacteria (Fig. 2F). Thus a flux of lactose, lactate, but also 1,2-PD that is only produced in presence of 2’-FL, was required for sustaining butyrate producing bacteria in our simulations.

We next turned to the model with the simplified consortium of species, the two *Bifidobacterium* subspecies and three butyrate producing species, to test if uptake of lactose, lactate and 1,2-PD was also required for butyrate producing bacteria to become abundant with this consortium. After blocking the uptake of lactose, lactate, or 1,2-PD by butyrate producing bacteria, the abundance of butyrate producing bacteria was reduced at the end of the simulations compared to the control (Fig. 3A). Surprisingly, however, and in contrast to the complete system (Fig. 2F), butyrate producing populations retained an abundance over 1 · 10^10^ bacteria in respectively 27 and 30 of 30 simulations when lactose or 1,2-PD uptake was disabled. Thus neither lactose nor 1,2-PD were essential for butyrate producing bacteria. Altogether, 1,2-PD, and thus 2’-FL, was required for butyrate producing bacteria in the complex system, but not in the simplified system. Thus these model simulations suggest that supplementation with 2’-FL introduces a flux of 1,2-PD from *Bifidobacterium* spp. to butyrate producing bacteria that prevents competitive exclusion of butyrate producers by competitors such as *B. vulgatus* (fig. 2C) or *C. acnes* (fig. 2D).

**Figure 3:**
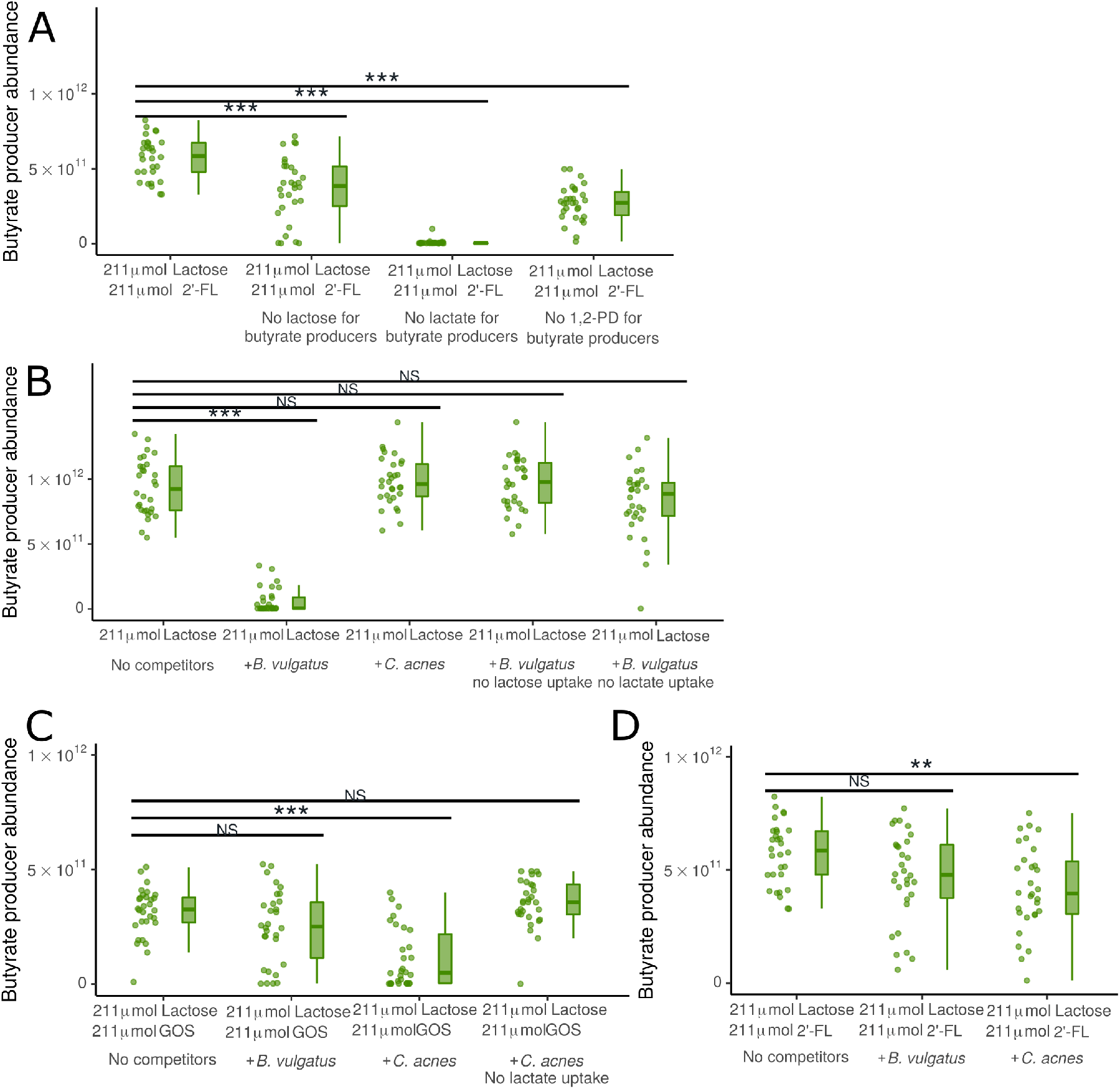
2’-FL makes butyrate producing bacteria resistant to competition by other infant gut bacteria. (A) Abundance of butyrate producers with 2’-FL and without competitors (only *Bifidobacterium* and butyrate producers) at the end of 21 days. Uptake of lactose, lactate, or 1,2-PD is disabled for butyrate producers in the ‘no lactose’, ‘no lactate’, and ‘no 1,2-PD’ conditions respectively. n=30 for each condition. Each simulation is represented by one dot. (p<0.001 for each disabled uptake compared to the baseline) (B,C,D) Abundance of butyrate producers at the end of 21 days (B) without prebiotics, either without competitors (only *Bifidobacterium* and butyrate producers), with addition of *B. vulgatus*, with addition of *B. vulgatus* unable to take up either lactose or lactate, or with addition of *C. acnes*. n=30 for each condition. Each simulation is represented by one dot. p<0.001 for abundance of butyrate producers with *B. vulgatus* compared to no competitors (C) with GOS, either without competitors (only *Bifidobacterium* and butyrate producers), with addition of *C. acnes*, with addition of *C. acnes* unable to take up lactate, or with addition of *B. vulgatus*. n=30 for each condition. Each simulation is represented by one dot. p<0.001 for abundance of butyrate producers with *C. acnes* compared to no competitors (D) with 2’-FL, either without competitors (only *Bifidobacterium* and butyrate producers), with addition of *C. acnes*, or with addition of *B. vulgatus*. n=30 for each condition. Each simulation is represented by one dot. p=0.001 for abundance of butyrate producers with *C. acnes* compared to no competitors. NS: Not significant, *: p<0.05, **:p<0.01, ***:p<0.001

### 2.4 *Bacteroides vulgatus* and *C. acnes* are effective competitors on different substrates

In the 2’-FL condition butyrate producing bacteria fed on lactate and 1,2-PD (Fig. 2E). In the conditions without 2’-FL no 1,2-PD was produced and lactate was consumed by *B. vulgatus* or *C. acnes* (Fig. 2C&D). This suggests that, in absence of 1,2-PD, *B. vulgatus* and *C. acnes* outcompete the butyrate producing bacteria for lactate. To investigate whether these species could indeed be responsible for outcompeting butyrate producing bacteria we again turned to the model with the simplified consortium and added the potential competitors *B. vulgatus* and *C. acnes* to the consortium one by one.

First we studied the simplified consortium in absence of prebiotics in the conditions with 211 µmol and 422 µmol lactose per three hours. The abundance of butyrate producing bacteria was reduced in presence of *B. vulgatus* but not in presence of *C. acnes* (Fig. 3B, 422 µmol visualized in Fig. S3). After blocking lactose or lactate uptake by *B. vulgatus* in the condition with 211 µmol lactose, the abundance of butyrate producing bacteria was restored (Fig. 3B), indicating that *B. vulgatus* required both lactose and lactate to effectively outcompete the butyrate producing bacteria.

In the conditions with GOS, the situation was reversed: *C. acnes* but not *B. vulgatus* outcompeted butyrate producing bacteria (Fig. 3C). After blocking uptake of lactate by *C. acnes* the abundance of butyrate producing bacteria was restored (Fig. 3C). *C. acnes* does not use lactose in the model. Taken together, these simulations suggest that lactate is required for competitive exclusion of butyrate producing bacteria by *C. acnes*.

In the condition with 2’-FL *B. vulgatus* did not outcompete butyrate producing bacteria (Fig. 3D). *C. acnes* (p=0.001) moderately suppressed butyrate producing bacteria, with 29 of 30 simulations still predicting an abundance of butyrate producing bacteria of over 1 ·10^10^ bacteria. This agrees with the simulations using the full consortium (Fig. 2B), which also displayed a robust abundance of butyrate producing bacteria in the 2’-FL condition.

### 2.5 Butyrate producing bacteria can use a mixture of lactate and 1,2-PD as substrates in the 2’-FL condition to grow faster than their competitors

To analyse how butyrate producing bacteria can outcompete other species only in the presence of 2’-FL but not in the presence of GOS or without prebiotics, we next examined the growth rates per timestep on unlimited quantities of the three key substrates for butyrate producing bacteria indicated above: lactose, lactate, and 1,2-PD. With unlimited availability of lactose, the growth of the three butyrate producing species was reduced relative to the growth of most other species (Fig. 4A). With unlimited lactate, growth for butyrate producing species was superior to the other species, but not to *C. acnes* (Fig. 4B). In presence of unlimited 1,2-PD and acetate the butyrate producing species *A. hallii* and *Roseburia inulinivorans* grew faster than the other species (Fig. 4C). On a mixture of limited lactate and 1,2-PD, with acetate available, two of the three butyrate producing species also grew faster compared to all other species (Fig. 4D). Thus the unique ability of butyrate producing bacteria to grow on 1,2-PD and acetate in the model allowed them to outcompete other lactate-consuming species in environments with 1,2-PD, such as those where *Bifidobacterium* consumes 2’-FL. However, they would be unable to outcompete the same species in conditions without 1,2-PD.

**Figure 4:**
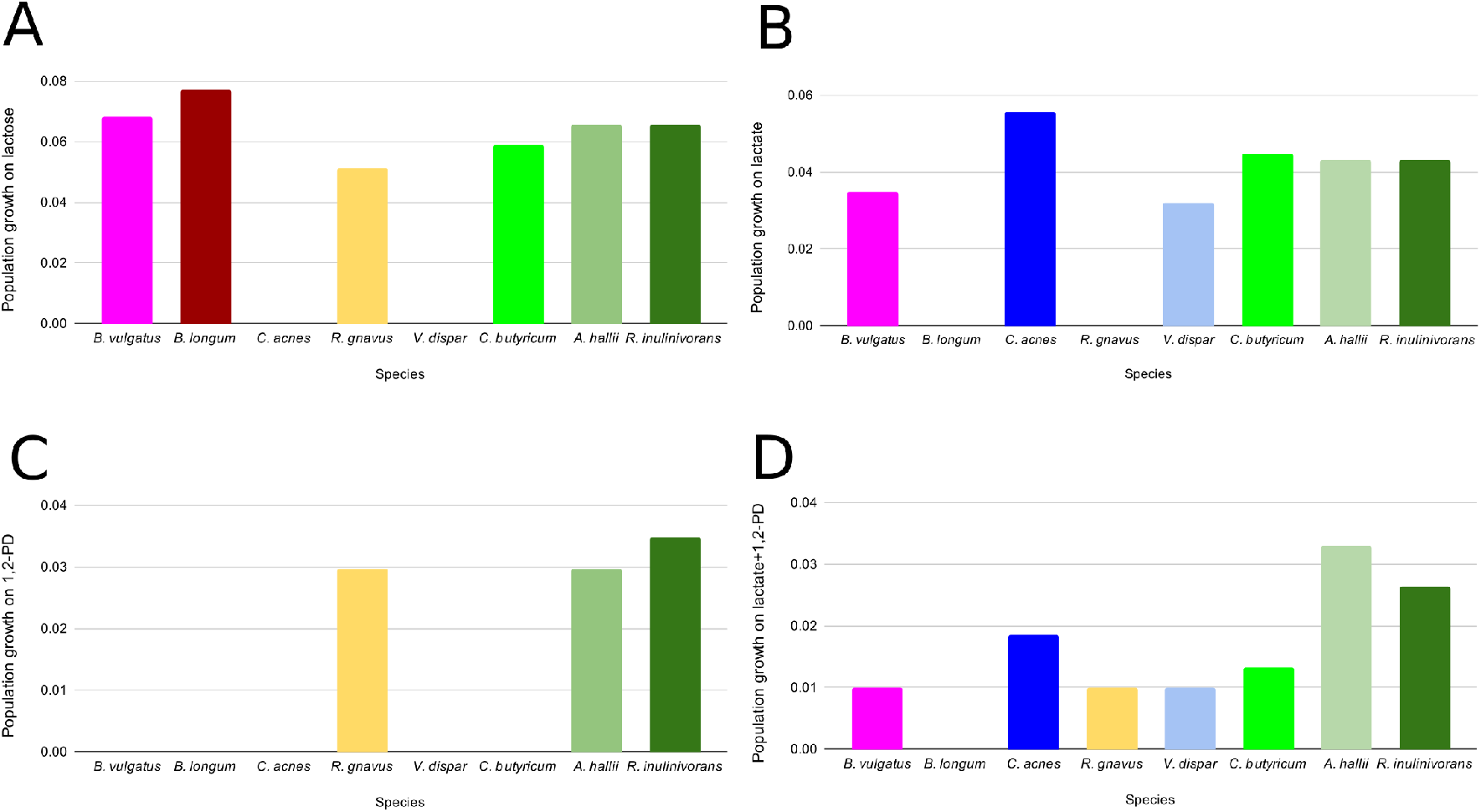
Populations of butyrate producing bacteria only grow much faster than their competitors on a mixed substrate of 1,2-PD and lactate. (A) Growth on unlimited lactose and water over a single timestep for butyrate producing bacteria (three rightmost bars, in green) compared to other lactose-fermenting bacteria in the model. (B) Growth on unlimited lactate and water over a single timestep for butyrate producing bacteria (three rightmost bars, in green) compared to other lactate-fermenting bacteria in the model. (C) Growth on unlimited 1,2-PD, acetate, and water over a single timestep for butyrate producing bacteria (two rightmost bars, in green) compared to another 1,2-PD-fermenting bacterial species in the model. (D) Growth on 1 µmol per ml of 1,2-PD and lactate, and unlimited acetate and water, over a single timestep for butyrate producing bacteria (three rightmost bars, in green) compared to other bacteria in the model for populations of 5 · 10^9^ bacteria with access to one lattice site (0.05ml)

### 2.6 Sensitivity analysis

Finally, to test the generality of our observations we performed a sensitivity analysis on the system. The enzymatic constraint (Fig. S2A&B), the death rate and growth rate (through the ATP required per population unit) (Fig. S2C&D), the placement of new populations of random species in empty lattice sites (colonization) (Fig. S2E&F), the diffusion of metabolites and populations (Fig. S2G&H),the amount of initial oxygen (Fig. S2I&J), and quiescence for large populations (Fig. S2K) were varied. We used three conditions for most changed parameters: 211 µmol lactose, 211 µmol lactose plus 211 µmol GOS, and 211 µmol lactose plus 211 µmol 2’-FL per three hours. We only used the latter two for disabling quiescence, as no populations entered quiescence during our initial runs with 211 µmol lactose. We found minor sensitivity for most parameter changes (Fig. S2). We found the most notable effects when we disabled colonization or initial oxygen. When we disabled colonization the abundance of butyrate producing bacteria was lower in all three conditions (p<0.001 for all). The absence of initial oxygen increased the abundance of butyrate producing bacteria in the condition without prebiotics and with 2’-FL (p=0.002,p=0.035). This indicates that the presence of initial oxygen and sustained colonization are particularly important in the simulated system.

## 3 Discussion

This paper describes a computational study of the effects of the prebiotics GOS and 2’-FL on butyrate producing bacteria in the infant gut microbiota. We have used the model to generate novel hypotheses to explain the — sometimes counter-intuitive — mechanisms at the biochemical and population level that underlie the effects of prebiotics. The model predicts that butyrate producing bacteria can coexist with *Bifidobacterium* in the infant gut with or without GOS or 2’-FL as long as no other bacterial species are present. As soon as other bacterial species are introduced into the model, we found that they can act as competitors, thus reducing the abundance of butyrate producing bacteria. Specifically, the model predicts that *B. vulgatus* outcompetes butyrate producing bacteria in absence of prebiotics. The predicted mechanism is that *B. vulgatus* consumes lactose and lactate, important food sources of the butyrate producing species. In presence of GOS, the model predicts that *C. acnes* becomes the key competitor of the butyrate producing bacteria due to its lactate consumption. In presence of 2’-FL, however, the butyrate producing species are no longer outcompeted. The mechanism as predicted by the model is as follows. The breakdown of 2’-FL by *Bifidobacterium* produces 1,2-PD. 1,2-PD becomes an additional food source for the butyrate producing bacteria, helping them to outgrow competitors. Thus, our modeling results predict that only 2’-FL, but not GOS supports populations of butyrate producing bacteria in their competition against species such as *B. vulgatus* and *C. acnes*.

The following *in vitro* and *in vivo* observations agree with these model predictions. Firstly, the model predicts co-existence and crossfeeding between *Bifidobacterium* and butyrate producing species on 2’-FL. In agreement with the model predictions, co-existence of and cross-feeding between *Bifidobacterium* and butyrate producing bacteria occurs *in vitro* within simplified, synthetic communities on glucose, fucose, and 2’-FL in the absence of competitors [64]. Secondly, the model predicts that in presence of the competitors such as *B. vulgatus* and *C. acnes*, *B. vulgatus* will become abundant in absence of prebiotics and outcompete butyrate producing species. In agreement with this model prediction, *B. vulgatus* is often abundant in the *in vivo* infant gut microbiota [4], and it can consume lactose *in vitro* [66]. No information is available on lactate consumption of *B. vulgatus*, but the related *Bacteroides fragilis* is able to consume lactate *in vitro* [42]. Thirdly, the model predicts that *C. acnes* outcompetes butyrate producing bacteria in presence of GOS by consuming lactate. In agreement with this prediction, *C. acnes* is found in 22% of infants [4] and *Cutibacterium avidum*, closely related to *C. acnes* [62], reduces the abundance of the butyrate producer *A. hallii* in an *in vitro* lactate-fed microbiota from infant fecal samples [56]. Both *C. acnes* and *C. avidum* consume lactate *in vitro* [29]. Finally, the model predicts that butyrate producing bacteria become competitive through cross-feeding on 1,2-PD, which is produced by *Bifidobacterium longum* from 2’-FL. In agreement with this prediction, the butyrate producer *A. hallii* cross-feeds on 1,2-PD in an *in vitro* synthetic community of *A. hallii* and *B. longum* [64]. Also in line with this prediction, 2’-FL supplementation increased the abundance of butyrate producing bacteria in *in vitro* fecal communities based on infant fecal samples, which likely include key competitors of butyrate producing species [73]. An *in vitro* colonic fermentation model inoculated with infant feces has previously been used to study the effect of introducing specific competitors to a lactate-consuming infant gut microbiota [56]. This approach could also be used to test if *B. vulgatus* and *C. acnes* are viable competitors in the infant gut and if the presence of 1,2-PD allows butyrate producing species to outcompete other bacteria.

More broadly, the model simulations without prebiotics predict that *Escherichia*, *Bacteroides*, and *Bifidobacterium* become the three most abundant genera, which agrees with the most abundant genera found in the infant gut microbiota around the age of three weeks [4, 69]. The relative abundances the model predicts for butyrate producing species range from 1.4% without prebiotics to 4.8% with 2’-FL, both of which are within the broad range of values reported for the butyrate producing community [3]. However, for two less abundant groups, Bacilli and *Veillonella*, the model predictions disagree with *in vivo* data. Firstly, an initially dominant Bacilli phase is sometimes seen *in vivo*, e.g. in 17.6% of subjects in [19], but not in any model outcomes. An initially dominant Bacilli phase is associated in non-premature infants with a shorter gestational period [19], but it is unclear exactly what factors are responsible. A similar initial dominance of Bacilli that often occurs in premature infants has been hypothesised to be related to selection pressures by the immune system, a different composition of initial colonizers [39], or a defective mucin barrier [18]. Secondly, the model predicted a very low *Veillonella dispar* abundance in all conditions. These predictions contradict *in vivo* data [55, 4] in which *V. dispar* is relatively abundant. *V. dispar* likely has a lower abundance in the model due to an incorrectly reduced growth rate relative to the other species in the model on lactate, the primary energy source of *V. dispar* [60], lactate, (Fig. 4B). We do not expect a large influence on the overall model predictions from this discrepancy, as *C. acnes* has a metabolism similar to that of *V. dispar* in the model and *in vitro*: both produce propionate, consume lactate, and cannot consume lactose [29]. However, we cannot exclude that other species in the model, such as *Veillonella* spp., may be more important competitors *in vivo* than the competitors that the model predicts.

Potential sources of the discrepancies between model predictions and experimental data include: (1) errors in the metabolic predictions of the underlying FBA models; (2) computational errors, and (3) incomplete representation of the biology underlying infant digestion. A typical error occurring in FBA models is an incomplete prediction of metabolic shifts, which is in part due to the assumption of FBA models that the growth rate or energy production is optimised [52]. For example, the FBA model does not correctly predict the metabolic shift from high-yield to low-yield metabolism as observed *in vitro* in *Bifidobacterium* growing on increasing concentrations of GOS and 2’-FL [84, 16]. FBA only predicts high-yield metabolism. The model, therefore, likely underestimates total lactate production. The effects of this discrepancy on the results are difficult to predict, but as lactate is a cross-feeding substrate, the underestimation of lactate may cause the model to underestimate the abundance of cross-feeding species such as *C. acnes* or butyrate producing bacteria. The optimality assumption of FBA also ignores any other ‘task’ that a bacterium has, besides growth. For example, sporulation, toxin production, or metabolic anticipation [48] may limit biomass production. The model does not represent such genetically regulated mechanisms.

Further errors in the model predictions can be due to simplifications in the FBA model. For example, we assume that the total flux through the reaction networks is capped (Eq. 4), so as to mimic the maximum volume in a cell that can be filled with enzymes. Here each enzyme is assumed to have equal maximum flux, and the optimization problem then predicts the optimal relative flux distribution. In reality, due to differences in enzyme concentration and enzyme efficiency these maximum fluxes can of course differ, which affects the predictions of FBA [6, 74]. If species-specific data on efficiency and genetic regulation of pathways become available, such weighting could be included in the model. The metabolic predictions from the FBA layer could be further improved in future versions of the model by integrating thermodynamic plausibility and favorability into FBA, which have previously improved metabolic predictions for intracellular metabolism [34, 25]. Additionally, the FBA model includes an extracellular compartment in which long GOS chains are broken down to shorter GOS chains, but it is not possible for extracellular breakdown products to diffuse during this process. Such extracellular digestion may lead to additional competition effects, because competitors may ‘steal’ digestion products without investing in the enzymes themselves [30]. Such effects may become important if additional species are introduced in the model that digest prebiotics extracellularly, such as *Bifidobacterium bifidum* [8].

Computational errors in the model (2) include the discretization of time, the discretization of space, and rounding errors in the FBA solver. Firstly, all processes in the model are assumed to be constant within each timestep, which means the model only roughly approximates the continuous temporal dynamics of processes such as metabolism and diffusion. Secondly, we discretize the three-dimensional continuous cylindrical space of the gut into a two-dimensional rectangular grid of lattice sites. We consider each lattice site to be of equal volume and to have equal flow through it. This simplification introduces many errors, as lattice sites must represent different shapes of three-dimensional space, and these shapes are not connected as they would be in three-dimensional space. It is difficult to estimate what impact these discretizations have on the model. Finally, the FBA solver uses floating point arithmetic to generate a deterministic but not exact solution to each FBA problem. These distortions are very small, typically on the order of 10^−15^ µmol per metabolite per FBA solution, so we do not expect a notable effect on the results.

Errors in the model predictions due to an incomplete representation of the biology underlying infant digestion (3) include missing organisms, missing ecological interactions, the simplifications we made to the metabolic input, and missing representation of host interactions. Firstly, the model does not include fungi or archaea in the infant gut. Both groups occur at a lower absolute abundance than the bacterial microbiota, but may still influence it [59]. Secondly, the model does not include interactions between bacteria other than cross-feeding and competition for resources. Missing interactions include acidification of the gut [14], the production of bacteriocins [22] and the effects of phage infections [47], all of which have species-specific effects. Thirdly, the model does not include the input of fats, proteins, or minerals into the gut. This means that the model cannot represent stimulation of growth by digestion of fats or proteins, nor potential limits on growth due to, for example, the lack of iron [46] or essential amino acids [41]. Finally, the model does not represent the interactions of the host with the microbiota, such as the continuous secretion by the gut wall of mucin [37] and oxygen [1], and the uptake of short-chain fatty acids [79]. Colonic mucins in particular could greatly influence the microbiota, as *B. bifidum* consumes colonic mucins extracellularly, which facilitates cross-feeding by butyrate producing bacteria *in vitro* [10].

Despite the inevitable limitations of the model, we have shown here how the model can be used to produce testable predictions on the effects of prebiotics and competition on butyrate producing bacteria in the infant gut microbiota. Future versions of the model may be a useful help in follow-up studies on the effects of nutrition on bacterial population dynamics in the infant and adult gut microbiota.

## 4 Methods

We used a spatially explicit model to represent the newborn infant gut microbiota. The model is based on our earlier models of a general microbiota [75] and the infant microbiota [78]. Prebiotic digestion is the most important addition in the present version of the model.

The model consists of a regular square lattice of 225 × 8 lattice sites, with each lattice site representing 2 × 2 mm of space. Taken together this represents an infant colon of 450 × 16 mm, in line with *in vivo* estimates [72, 15]. Each lattice site can contain an amount of the 735 metabolites represented in the model, as well as a single bacterial population.

### 4.1 Species Composition

Species were selected based on [4], using sheet 2 of their Table S3. We selected the 20 entries with the highest prevalence in vaginally delivered newborns. After removing two duplicate entries we selected a representative species for each genus from the AGORA database [43]. We then added an additional *Bifidobacterium longum* ssp. *infantis* GEM to serve as prebiotic consumer, and a *Roseburia inulinivorans* GEM. *Roseburia* spp. have been shown to be a prevalent butyrate producing bacterium in infants in other studies [3]. Together, these form the list of species (Table 1).

### 4.2 Changes from AGORA

The model uses GEMs generated in the AGORA project [44]. We have applied various changes and additions to these models (Table S1).

We have added digestion of GOS or 2’-FL to the *B. longum* ssp. *infantis* GEM as follows. 2’-FL digestion was implemented by adding reactions representing an ABC-transporter and an intracellular fucosidase that breaks 2’-FL down to lactose and fucose [84]. GOS was represented through separate DP3, DP4, and DP5 fractions [77]. The DP4 and DP5 fractions are broken down extracellularly to DP3 and DP4 fractions respectively, releasing one galactose molecule in the process [76]. The DP3 fraction is taken up with an ABC transporter, and broken down internally to lactose and galactose [76].

We have also further expanded earlier curation of the AGORA GEMs [78]. We disabled anaerobic L-lactate uptake for the *Bifidobacterium* GEMs and for *E. coli* in line with available literature [23, 13]. To have the GEMs correspond with existing literature on lactose uptake we added a lactose symporter to *Anaerobutyricum hallii* [10], both *Bifidobacterium longum* GEMs [54], *Roseburia inulinivorans* [57], *Haemophilus parainfluenzae* [32], and *Rothia mucilaginosa* [71]. We also added galactose metabolism to *R. inulinivorans* [35] and *R. mucilaginosa* [71]. Further changes were made to prevent unrealistic growths and the destruction of atoms within reactions (Table S1).

### 4.3 Validity checks

After applying the changes in Table S1 we tested all GEMs individually for growth on a substrate of lactose and water. In line with literature, this did not lead to growth for *Veillonella disparans* [60], *Cutibacterium acnes* [29], *Eggerthella* sp. YY7918 [83], and *Gemella morbillorum* [80]. All other species grew on this substrate. We also checked for any spurious growth by checking each GEM for growth with only water present.

During each simulation, the model checks the FBA solutions for thermodynamic plausibility. The model uses a database of Gibbs free energy values [50] for all metabolites except 2’-FL and GOS. Values for 2’-FL and GOS were generated from the values for lactose and fucose, and lactose and galactose, respectively. Separate values were generated for the separate fractions of GOS. All values assumed a pH of 7 and an ionic strength of 0.1 M. We found that in the simulations of Fig. 2A with the baseline level of lactose, combined with those with GOS and 2’-FL (n=90) 99.98% of all FBA solutions had a lower or equal amount of Gibbs free energy in the output compared to the input. The remaining 0.02% of FBA solutions was responsible for 0.003% of total bacterial growth.

### 4.4 FBA with enzymatic constraint

Although other aspects of the model were changed, the FBA approach we used is identical to that used in the earlier model [78]. The model uses a modified version of flux balance analysis with an enzymatic constraint to calculate the metabolic inputs and outputs of each population at each timestep [52, 45]. Each GEM is first converted to a stoichiometric matrix 𝑆. Reversible reactions are converted to two irreversible reactions, so that flux is always greater than or equal to 0. Reactions identified in the GEM as ‘exchange’, ‘sink’, or ‘demand’ in the GEM are also recorded as ‘exchange’ reactions. These exchange reactions are allowed to take up or deposit metabolites into the environment. Each timestep, all reactions are assumed to be in internal steady state:

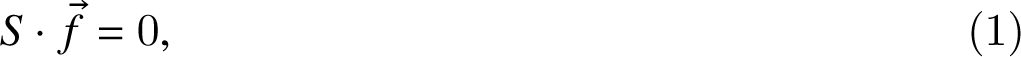

where 𝑓^➔^ is a vector of the metabolic fluxes through each reaction in the network, in mol per time unit per population unit.

Each exchange reaction that takes up metabolites from the environment 𝐹_𝑖𝑛_is constrained by an upper bound 𝐹_𝑢𝑏_which represents the availability of metabolites from the environment. It is determined as follows:

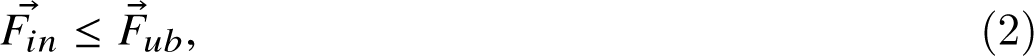

where 𝐹^➔^_𝑖𝑛_ is a vector of fluxes between the environment and the bacterial population. 𝐹^➔^_𝑢𝑏_ is a vector of upper bounds on these fluxes. 𝐹^➔^_𝑢𝑏_ is set dynamically at each timestep 𝑡 by the spatial environment at each lattice site 𝑥➔:

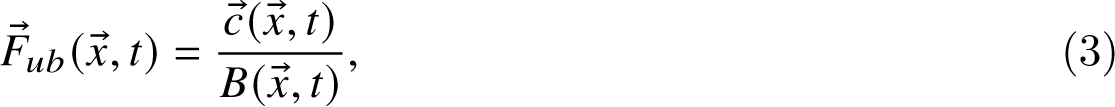

where 𝑐➔ is a vector of all metabolite concentrations in mol per lattice site, 𝑥➔ is the location and 𝐵(𝑥➔, 𝑡) is the size of the local bacterial population. The size of 𝐵 can range from 5 · 10^7^ to 2 · 10^10^ bacterial cells.

Finally the enzymatic constraint constrains the total flux through the network. It represents the maximum, total amount of flux that can be performed per cell in each population:

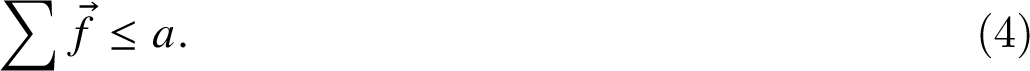

The enzymatic constraint 𝑎 is in mol per time unit per population unit. As both 𝑓^➔^ and 𝑎 are per population unit, this limit scales with population size, so each bacterial cell contributes equally to the metabolic flux possible in a lattice site. The enzymatic constraint is included as a constraint on each FBA solution. Given the constraints, FBA identifies the solution that optimises the objective function, ATP production. The solution consists of a set of input and output exchange fluxes 𝐹^➔^_𝑖𝑛_ (𝑥➔, 𝑡) and 𝐹_𝑜_^➔^_𝑢𝑡_ (𝑥➔, 𝑡), and a growth rate 𝑔(𝑥➔, 𝑡). The exchange fluxes are taken as the derivatives of a set of partial-differential equations to model the exchange of metabolites with the environment. The size of the population increases proportionally to the growth rate in the FBA solution.

To mimic quiescence at high densities, populations above the spreading threshold of 2 · 10^10^ bacteria do not perform metabolism. In practice this rarely occurs because we maintain sufficient space for populations to spread into empty lattice sites. In the simulations of Fig. 2A (n=120) metabolism was not performed in, on average, 0.05% of all populations in a timestep.

### 4.5 Environmental metabolites

We model 735 different extracellular metabolites. This is the union of all metabolites that can be exchanged with the environment by at least one GEM in the model. In the simulations 39 metabolites are present in the medium in more than micromolar amounts at any point. We combine the L-lactate and D-lactate metabolites for fig. 1B, Video S1 and Video S2. Nearly all lactate in the model is L-lactate.

To represent the mixing of metabolites by colonic contractions we apply a diffusion process to the metabolites at each timestep. Metabolic diffusion is applied in two equal steps to the model. In each step, 14.25% of each metabolite diffuses from each lattice site to each of the four nearest neighbours. This causes a net diffusion each timestep of 6.3 · 10^5^ 𝑐𝑚^2^/s. Metabolites are also added and removed by bacterial populations as a result of the FBA solutions, yielding

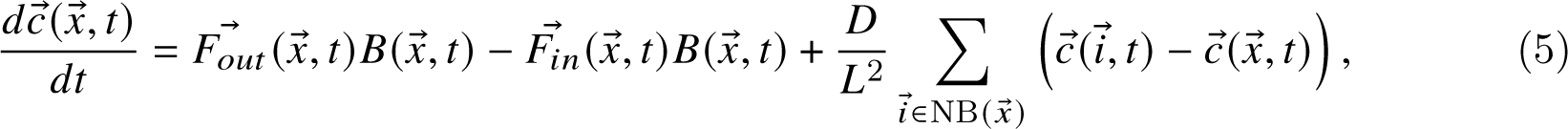

where 𝐹_𝑜_^➔^_𝑢𝑡_ (𝑥➔, 𝑡) is a vector of fluxes from the bacterial populations to the environment, in mol per time unit per population unit, 𝐷 is the diffusion constant, 𝐿 is the lattice side length, and 𝑁 𝐵(𝑥➔) are the four nearest neighbours.

All metabolites except oxygen are moved distally by one lattice site every timestep to represent advection. The transit time, including diffusion, is approximately 11 hours, corresponding with *in vivo* observations in newborn infants [61, 36]. Metabolites at the most distal column of the lattice, the end of the colon, are removed from the system at each timestep.

Every 60 timesteps (representing three hours) metabolites representing inflow from the small intestine are inserted into the first six columns of lattice sites. Three hours represents a realistic feeding interval for neonates [31]. Food intake contains 211 µmol of lactose by default, a concentration in line with human milk [5], assuming 98% host uptake of carbohydrates before reaching the colon [12]. In some simulations 211 µmol of additional lactose, GOS, or 2’-FL is added. Because there is very little uptake of prebiotics by the infant [27], the oral intake of prebiotics would be much lower than that of lactose. GOS is inserted as separate fractions of DP3, DP4, or DP5 based on analysis of the composition of Vivinal-GOS [77]. 64% is DP3, 28% is DP4 and 8% is DP5. Water is provided in unlimited quantities. Oxygen is placed during initialisation [68] at 0.1 µmol per lattice site. No other metabolites are available, other than those produced as a result of bacterial metabolism within the model.

### 4.6 Population dynamics

During initialization there is a probability of 0.3 for each lattice site to get a population of size 5 · 10^7^ of a random species (Table 1). Taken together, this averages around 540 populations, leading to a total initial bacterial load of 2.7 · 10^10^, in line with *in vivo* estimates [53] when we assume a uniform bacterial density and a total colon volume of 90 ml. In each timestep each local population solves the FBA problem based on its own GEM, the enzymatic constraint 𝑎, its current population size 𝐵(𝑥➔, 𝑡) and the local concentrations of metabolites 𝑐➔(𝑥➔, 𝑡), and applies the outcome to the environment (see above) and the growth rate 𝑔(𝑥➔, 𝑡) to its own population size, as follows:

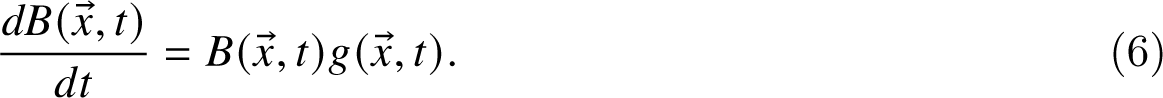

Each step, each population of at least 1 · 10^10^ bacteria (Table 2) will create a new population if an adjacent empty lattice site is available. Half of the old population size is transferred to the new population, so that the total size is preserved. To mimic colonisation new populations are introduced at random into empty lattice sites during the simulation, representing both dormant bacteria from colonic crypts [40] and small bacterial populations formed from ingested bacteria, which may only become active after being moved far into the gut. Each empty lattice site has a probability of 0.00005 (Table 2) each step to acquire a new population of a randomly selected species. All species have an equal probability to be selected. We initialise these populations at the same population size 𝐵 as the initial populations in the model (Table 2). Each population dies out at a probability of 0.0075 per timestep, creating a turnover within the range of estimated microbial turnover rates in the mouse microbiota [26].

To mix the bacterial populations the lattice sites swap population contents each timestep. We use an algorithm inspired by Kawasaki dynamics [38], also used previously for bacterial mixing [78, 75]: In random order, the bacterial content of each site, i.e., the bacterial population represented by its size 𝐵(𝑥➔, 𝑡) and the GEM, are swapped with a site randomly selected from the Moore neighbourhood. This swap only occurs if both the origin and destination site have not already swapped in this timestep. With this mixing method the diffusion constant of the bacterial populations is 6.3 · 10^5^𝑐𝑚^2^/𝑠, equal to that of the metabolites. Bacterial populations at the most distal column, i.e. at the exit of the colon, are removed from the system. To increase the bacterial diffusion rate in the sensitivity analysis this process was executed five times, marking all sites as unswapped after each execution. To decrease the bacterial diffusion rate the number of swaps was limited to a fifth of the usual number of swaps.

### 4.7 Analysis

We record the size, species, location, and important exchange fluxes 𝐹^➔^_𝑖𝑛_ (𝑥➔, 𝑡) and 𝐹_𝑜_^➔^_𝑢𝑡_ (𝑥➔, 𝑡) for each population at each timestep. To detect irregularities we also record the net flux of carbon, hydrogen, oxygen, and Gibbs free energy for every population at each timestep. Gibbs free energy is estimated using the Equillibrator database [50]. Energy loss 𝑙 in joules per timestep per population unit is recorded as follows, where 𝑖 are metabolites, 𝐹 is the exchange flux rate in mol per timestep per population unit and 𝐸 contains the Gibbs free energy in joules per mol for each metabolite,

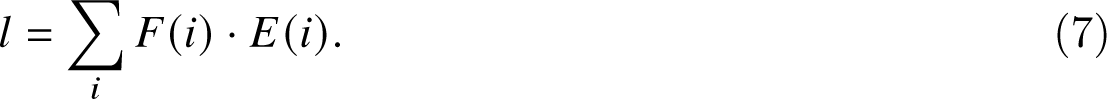

### 4.8 Parameters

Parameters of the system are listed in table 2. We estimate a total volume of 90ml for the infant colon [72, 15], which leads to a rough estimate on the order of 10^12^ bacteria in the newborn infant colon given an abundance per ml of around 10^10^ [53]. Values for free parameters were estimated and evaluated in the sensitivity analysis.

### 4.9 Implementation

We implemented the model in C++11. We based the model on our own earlier models of the gut microbiota [75, 78]. Random numbers are generated with Knuth’s subtractive random number generator algorithm. Diffusion of metabolites was implemented using the Forward Euler method. The GEMs are loaded using libSBML 5.18.0 for C++. We used the GNU Linear Programming Kit 4.65 (GLPK) as a linear programming tool to solve each FBA with enzymatic constraint. We used the May 2019 update of AGORA, the latest at time of writing, from the Virtual Metabolic Human Project website (vmh.life). We used Python 3.6 to extract thermodynamic data from the eQuilibrator API (December 2018 update) [50]. When not noted otherwise p-values were calculated with R 4.2.1 using the Mann-Whitney test from the ‘stats’ package 3.6.2. Model screenshots were made using the libpng16 and pngwriter libraries. Other visualisations were performed with R 4.2.1 and Google Sheets. Raincloud visualisations used a modified version of the Raincloud plots library for R [2].

## 5 Supplemental material

**S1 Table.**

**Table of changed or deleted reactions and annotations.csv**

A table of changes made to the AGORA models as a .csv file.

**S1 Video.**

**Video of a simulation with no prebiotics, consisting of a visualisation of the distribution of bacterial species and major metabolites.** Lines represent, from top to bottom: Bacteria, lactose, 2’-FL, lactate (Both L and D), acetate, 1,2-PD, butyrate, succinate, CO2, H2, propionate

**S2 Video.**

**Video of a simulation with 2’-FL, consisting of a visualisation of the distribution of bacterial species and major metabolites.** Lines represent, from top to bottom: Bacteria, lactose, 2’-FL, lactate (Both L and D), acetate, 1,2-PD, butyrate, succinate, CO2, H2, propionate

**S3 Video.**

**Video of a simulation without prebiotics, displaying fluxes between population and metabolite pools** Line width is scaled with the flux per metabolite over 60 timesteps per frame, multiplied by the carbon content of the molecule, with a minimum threshold of 100 µmol atomic carbon.

**S4 Video.**

**Video of a simulation with 2’-FL, displaying fluxes between population and metabolite pools.** Line width is scaled with the flux per metabolite over the 60 timesteps per frame, multiplied by the carbon content of the molecule, with a minimum threshold of 100 µmol atomic carbon.

## 6 Contributions

J.M.W.G., and R.M.H.M acquired funding. D.M.V., J.M.W.G., and R.M.H.M. conceived and planned the simulations. D.M.V. wrote software used for the simulations. D.M.V. performed the simulations and analyzed the data. R.S, E.L., J.M.W.G., and R.M.H.M contributed to the interpretation of the results. J.M.W.G., and R.M.H.M. supervised the project. D.M.V. drafted the manuscript. D.M.V., R.S., E.L., J.M.W.G. and R.M.H.M. revised and edited the manuscript.

## Acknowledgments

This study was financially supported by FrieslandCampina. R.S, E.L., and J.M.W.G. are currently employed by FrieslandCampina. This work was performed using the ALICE compute resources provided by Leiden University.

---

**S1 Figure.**
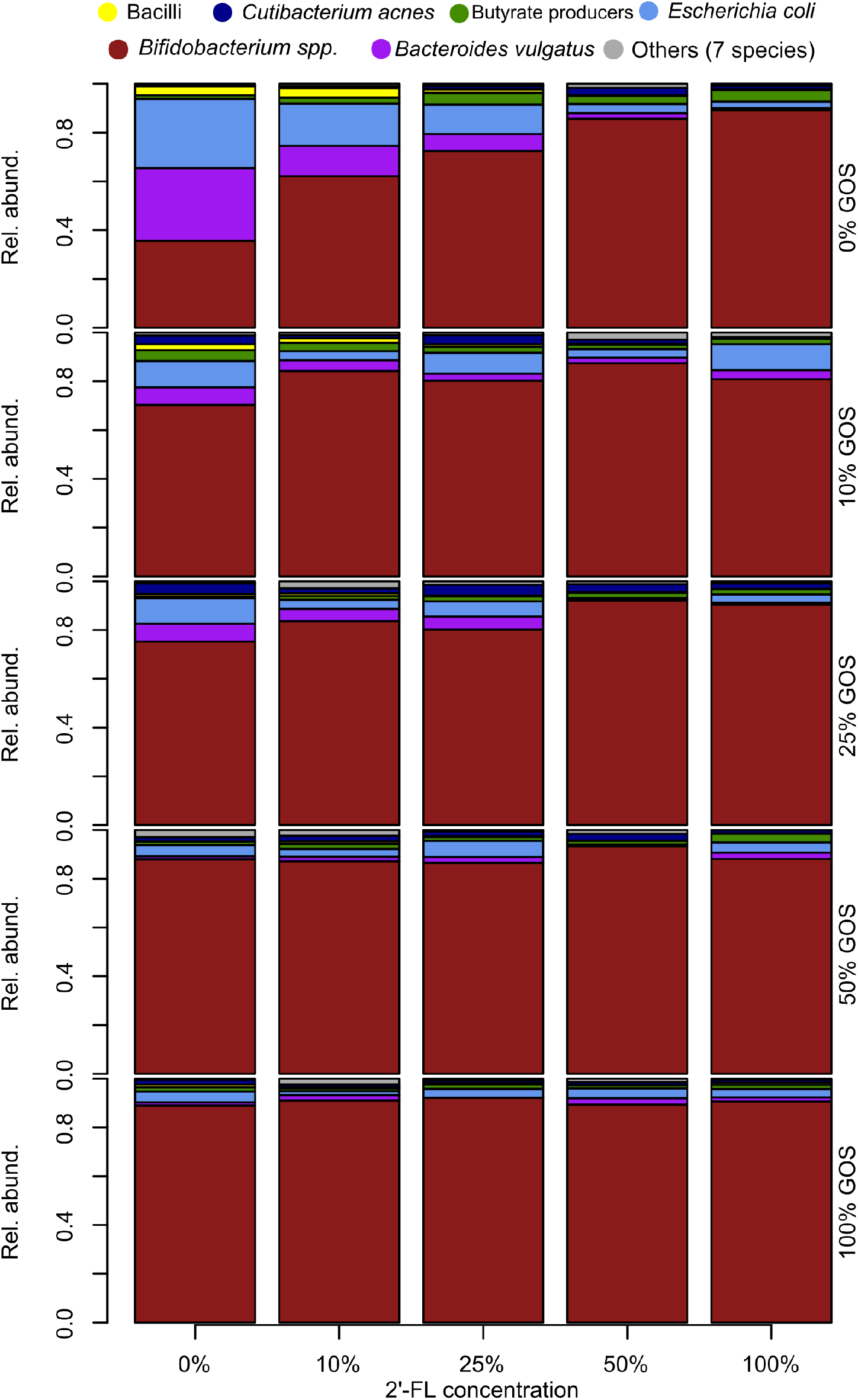
Relative abundance of bacterial species at the end of 21 days with varying inputs of 2’-FL and GOS compared to the fixed input amount of lactose. n=30 for each condition, each simulation is weighed equally.

**S2 Figure.**
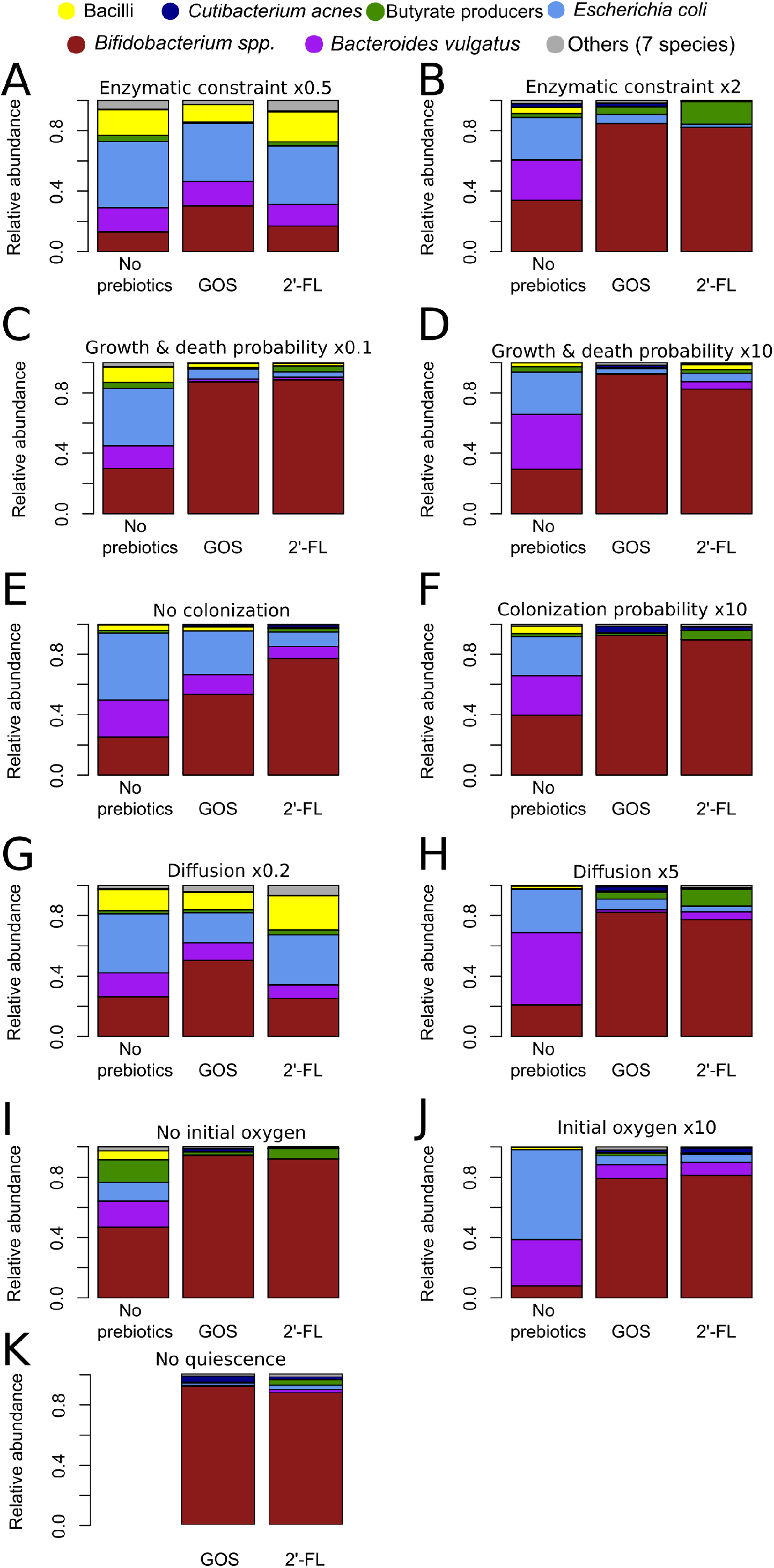
**(A to K) Relative abundance of bacterial species in the conditions with no pre-biotics, with GOS, or with 2’-FL at the end of 21 days**, with the following alteration from the baseline of Fig. 2A: (A) Enzymatic constraint loosened by a factor of 2, to 4 µmol flux per timestep per 1 · 10^10^ population (B) Enzymatic constrained tightened by a factor of 2, to 1 µmol flux per timestep per 1 · 10^10^ population (C) Growth decreased by a factor of 10, by increasing the ATP per bacteria to 1 · 10^−14^, with the death probability decreased to 0.00075 per population per timestep. (D) Growth increased by a factor of 10 by decreasing the ATP per bacteria to 1 · 10^−16^, with the death probability increased to 0.075 per population per timestep (E) Colonisation removed by setting the probability for new populations to be placed after initialization to 0 (F) Colonisation increased by x10 by setting the probability per empty lattice to acquire a new population to 0.0005 per timestep (G) Diffusion of both metabolites and bacteria decreased by a factor of 5 to 1.26 · 10^−6^ 𝑐𝑚^2^/s (H) Diffusion of both metabolites and bacteria increased by a factor of 5 to 3.15 · 10^−5^ 𝑐𝑚^2^/s (I) No initial presence of oxygen (J) Initial oxygen increased to 1 µmol per lattice site (K) Quiescence disabled For each figure: n=30 for each condition, each simulation is weighed equally.

**S3 Figure.**
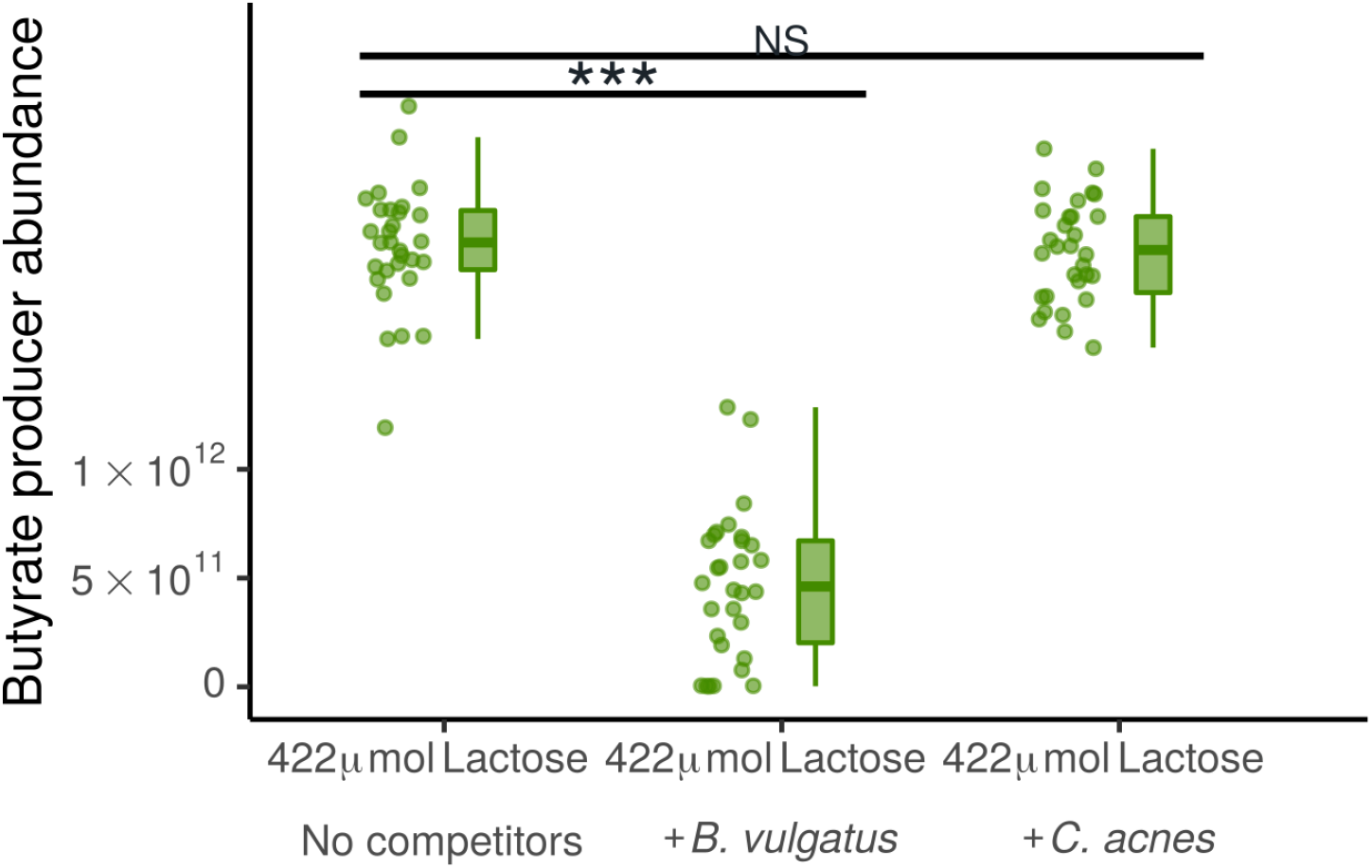
**Abundance of butyrate producing bacteria at the end of 21 days with 422 µmol of lactose per three hours and without prebiotics, either without competitors (only *Bifidobacterium* and butyrate producing bacteria), with addition of *B. vulgatus*, or with addition of *C. acnes***. n=30 for each condition. Each simulation is represented by one dot. NS: Not significant, *: p<0.05, **:p<0.01, ***:p<0.001

